# Evolutionary consequences of DNA methylation in a basal metazoan

**DOI:** 10.1101/043026

**Authors:** Groves Dixon, Line K. Bay, Mikhail V. Matz

**Affiliations:** Institute for Cell and Molecular Biology, University of Texas, Austin, USA; Australian Institute of Marine Science, PMB 3, Townsville, Queensland 4810, Australia; ARC Centre of Excellence for Coral Reef Studies, James Cook University, Townsville, Queensland 4811, Australia; Department of Integrative Biology, University of Texas, Austin, USA

## Abstract

Gene body methylation (gbM) is an ancestral and widespread feature in Eukarya, yet its adaptive value and evolutionary implications remain unresolved. The occurrence of gbM within protein coding sequences is particularly puzzling, because methylation causes cytosine hypermutability and hence is likely to produce deleterious amino acid substitutions. We investigate this enigma using an evolutionarily basal group of Metazoa, the stony corals (order Scleractinia, class Anthozoa, phylum Cnidaria). We show that gbM correlates with breadth and abundance of transcription and with slow sequence evolution. We also show a strong correlation between gbM and codon bias due to systematic replacement of CpG bearing codons. We suggest that the ancestral function of gene body methylation is tied to selective pressure for accurate and stable gene expression, and that mutation caused by gbM may be a previously unrecognized driver of adaptive codon evolution.

## Introduction

DNA methylation is an evolutionarily widespread epigenetic modification found in plants, animals and fungi. It is defined as the covalent addition of a methyl group to the one of the four DNA bases, predominantly on the fifth carbon of cytosines within CG dinucleotides (CpGs), producing 5-methylcytosine (5mC). Unlike other epigenetic modifications, DNA methylation not only alters chromatin structure and transcription, it changes the mutation rate of the underlying DNA. This is because 5mC undergoes deamination reactions more readily than normal cytosine (Shen *et al*. 1994) and deamination produces thymine rather than uracil, which is less likely to be accurately repaired (Zemach & Zilberman 2010). Because of this hypermutability, sequences that are heavily methylated in the germ-line become deficient in CpGs over evolutionary time, with corresponding increases in TpG and CpA dinucleotides (Sved & Bird 1990). Hence DNA methylation has evolutionary consequences outside of its direct physiological effects.

Evolutionary effects of 5mC hypermutability are apparent in both vertebrate and invertebrate genomes. In mammals, DNA methylation is ubiquitous, so that nearly all genomic regions show lower than expected frequency of CpGs (Karlin & Mrázek 1996; McGaughey *et al*. 2014). The exception is regions of elevated CpG content called CG islands that are protected from DNA methylation (Jones 2012). In most invertebrates, DNA methylation is not ubiquitous but patchy, with preferential occurrence on CpGs within gene bodies (Suzuki *et al*. 2007; Zemach *et al*. 2010). This form of DNA methylation, referred to as gene body methylation (gbM), is also found in mammals. Notably, one of the mammalian *de novo* DNA methyltransferases (DNMT3B1) recruits specifically to gene bodies, and binds preferentially to actively transcribed genes (Baubec *et al*. 2015), As gbM also occurs in plants (Tran *et al*. 2005; Zilberman *et al*. 2007) it likely represents an ancestral form of DNA methylation for eukaryotes (Feng *et al*. 2010).

Despite its widespread phylogenetic occurrence gbM is by no means universal. In several groups, including classical model organisms such as yeast (*Saccharomyces cerevisiae*), fruit fly (*Drosophila melanogaster*), and worm (*Caenorhabditis elegans*), DNA methylation is extremely scarce or lost altogether (Capuano *et al*. 2014). It has been proposed that the secondary loss of DNA methylation from these organisms occurred because its mutational costs outweighed its adaptive value (Zemach *et al*. 2010). Indeed even within gene bodies, methylation occurs preferentially on exons (Zemach *et al*. 2010; Wang *et al*. 2013), where mutations are likely to have the greatest deleterious effect. In humans, DNA methylation is a strong source of mutation (Cooper *et al*. 2010; Xia *et al*. 2012) and increases deleterious *de novo* mutations with paternal age (Francioli *et al*. 2015). Why, given its apparently nonessential and outright mutagenic nature, has gbM persisted for so long across such a diversity of taxa? Addressing this question requires understanding of both the adaptive value and evolutionary consequences of gbM in its ancestral form. To this end, we characterized the methylome of a basal metazoan—the reef-building coral *Acropora millepora*. Methylomic data were analyzed in the context of gene expression, substitution rates, and codon usage to elucidate functional and evolutionary impacts of gbM at the base of the metazoan tree.

## Results

### Using MBD-seq to quantify gene body methylation

We used Methylation Binding Domain enrichment sequencing (MBD-seq)(Harris *et al*. 2010) to measure gbM in *A. millepora*. The strength of methylation for 24320 coding regions was quantified as the log2 fold difference between captured and flow-through fractions of MBD enrichment preparations. We refer to this log2 fold difference as the MBD-score. Analysis of the distribution of MBD-scores (fig. 1A) showed that it was best described as a mixture of two or more Gaussian components (supplementary fig. S1). MBD-score correlated with CpGo/e, indicating that our measure of gbM overlapped closely with historical patterns of germ-line methylation (fig. 1B). As an MBD-score of zero indicated equal representation in the captured and flow-through fractions we chose this value to separate strongly and weakly methylated genes. Genes with MBD-scores greater than zero are referred to as strongly methylated genes, those with scores less than zero are referred to as weakly methylated.

**Figure 1:**
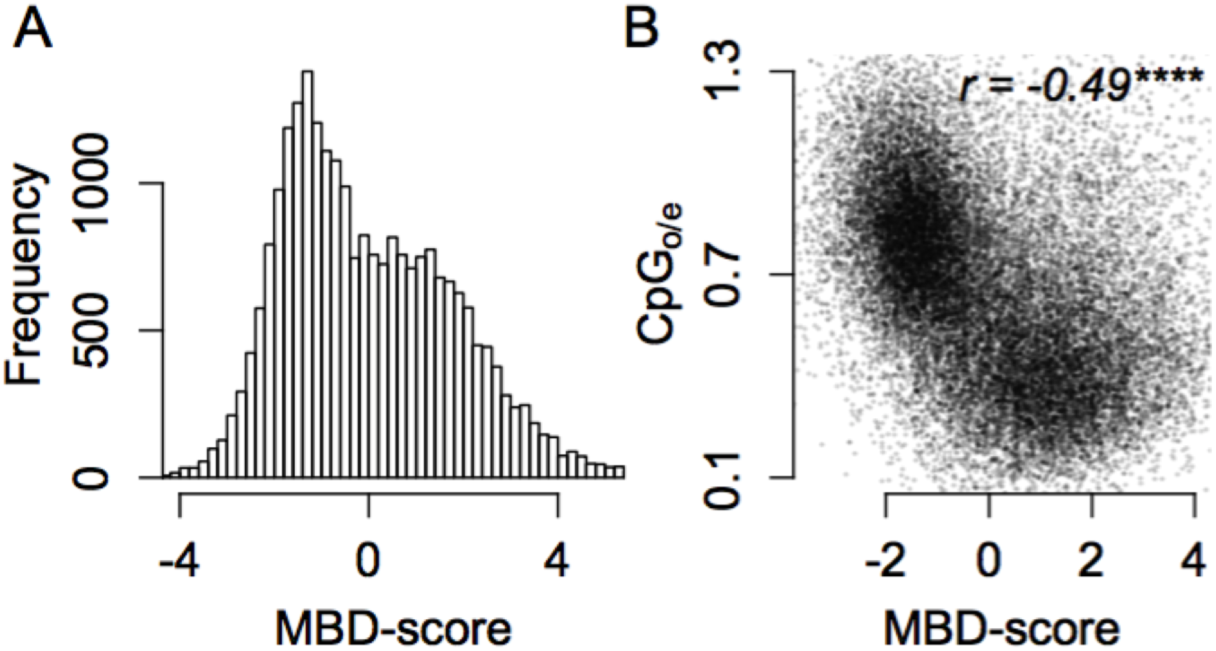
Methylation score is bimodally distributed and correlates with CpGo/e. (A) Distribution of MBD-score (log_2_ fold difference between enriched and flow-through MBD-seq libraries). Higher values indicate stronger methylation. (B) Scatterplot of MBD-score and CpGo/e. Lower values for CpGo/e are expected with stronger methylation. Asterisks indicate significance based on Spearman’s rho (ns > 0.05; * < 0.05; ** < 0.01; *** < 0.001; **** < 0.0001).

### MBD-score is linked with gene fonction and expression patterns

MBD-score correlated with gene function. Analysis of selected GO categories for biological processes revealed that strongly methylated genes tend toward biological functions that are spatially and temporally stable such as DNA metabolism, ribosome biogenesis, translation, RNA metabolism and transcription. Weakly methylated genes tended to involve biological functions that are spatially and temporally regulated, such as cell-cell signaling, response to stimulus, signal transduction, cell adhesion, defense response and development (supplementary fig. S2A). Clustering of KOG categories for higher or lower MBD-scores further supported these results (supplementary fig. S2B).

To directly examine the relationship between gbM and transcriptional stability we correlated MBD-score with RNA-seq data comparing different developmental stages and environmental conditions. For developmental stage, log2 fold differences in transcript abundance between *A. millepora* adults and larvae (described in Dixon *et al*. 2015) negatively correlated with MBD-score (fig. 2A). Significantly differentially expressed genes (DEGs at FDR < 0.01) were 1.4 times more frequent among weakly methylated genes (fig. 2B). A similar trend was found for variation in expression due to environmental conditions (fig. 2C-D). Here clonal fragments of adult colonies were exposed to two environmentally distinct regimes for three months prior to sampling for RNA-seq (Dixon *et al*. 2014). Differential expression (FDR < 0.01) between environmental regimes was 2.2 times more frequent among weakly methylated genes.

**Figure 2:**
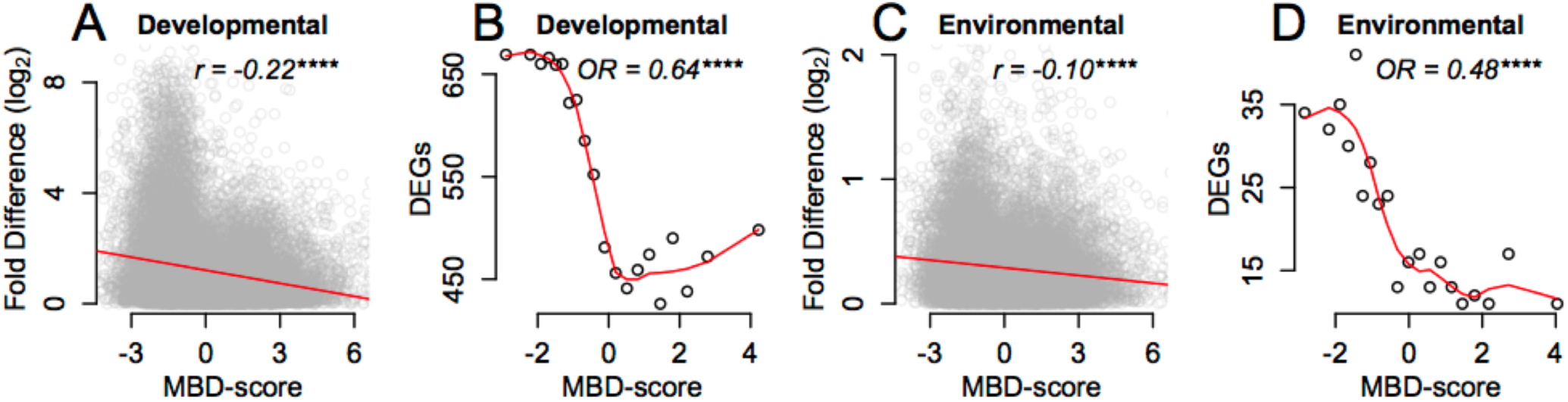
Gene body methylation predicts transcriptional stability across developmental stages and environmental regimes. (A) Scatterplot of MBD-score and transcriptional variation (given as log2 fold differences) between adult colonies and juvenile offspring. Red line shows least squared regression. Asterisks indicate significance based on Spearman’s rho. (B) Distribution of differentially expressed genes (DEGs; FDR < 0.01) between juveniles and adults. All genes were divided into 20 quantiles ranked by MBD-score. The number of differentially expressed genes in each quantile was plotted against the median MBD-score for that quantile. Enrichment of DEGs among the weakly methylated genes (MBD-score < 0) compared to strongly methylated genes (MBD-score > = 0) is given as the odds ratio (OR) for Fisher’s exact test. Red line shows a smoothed trace of the points fit with a span of 0.5. (C-D) The same figures representing transcriptional variation between populations of clonal colony fragments transplanted between distinct habitats as described in Dixon et al. (2014), Significance notation: ns > 0.05; * < 0.05; ** < 0.01; *** < 0.001; **** < 0.0001.

MBD-score also showed weak but significant correlation with transcript abundance (supplementary fig. S3A and B, Supplementary Material online). Highly expressed genes were on average the most strongly methylated (supplementary fig. S3C and D). The top 5% most strongly methylated genes however, showed lower average expression (supplementary fig. S3E). This indicates that while gbM is generally associated with elevated transcription, extreme levels may be inhibitory. This appears to be particularly true for short genes, as the removal of coding sequences shorter than 800 bp mitigated the trend (supplementary fig. S3F).

### Phylogeny

We used a conserved set of 192 coding sequences for phylogenetic construction. These sequences had >75% amino acid identity and 80% representation among the 20 species. Phylogenetic construction was performed using the GTRGAMMA model in RAxML (Stamatakis 2014). All bipartitions had 100% bootstrap support based on 1000 repetitions. All orders, families, and genera formed monophyletic groups (fig. 4). Species from the ‘complex’ and ‘robust’ coral clades (Romano & Palumbi 1996; Kitahara *et al*. 2010) also formed monophyletic groups. For the species in which they overlapped, our tree agreed fully with that published by Kitchen *et al*. (2015).

### Strongly methylated genes evolve slowly

Pairwise comparisons between orthologs from *A. millepora* and each of the other species revealed that strongly methylated genes evolve slowly. The trend was strongest for nonsynonymous substitutions (dN). When orthologs from *A. millepora* were compared with other *Acropora* species, mean dN was between 43% and 68% higher for weakly methylated genes than strongly methylated genes (fig. 3; supplementary fig. S4). Pairwise comparisons with all species outside of the *Acropora* genus produced similar results, with mean dN between 17% and 52% (mean = 36 ± SEM 3%) higher for weakly methylated genes (fig. 3; supplementary fig. S5). Negative correlation between dN and MBD-score was significant for all species comparisons (p << 0.001; Spearman’s Rank Test).

**Figure 3:**
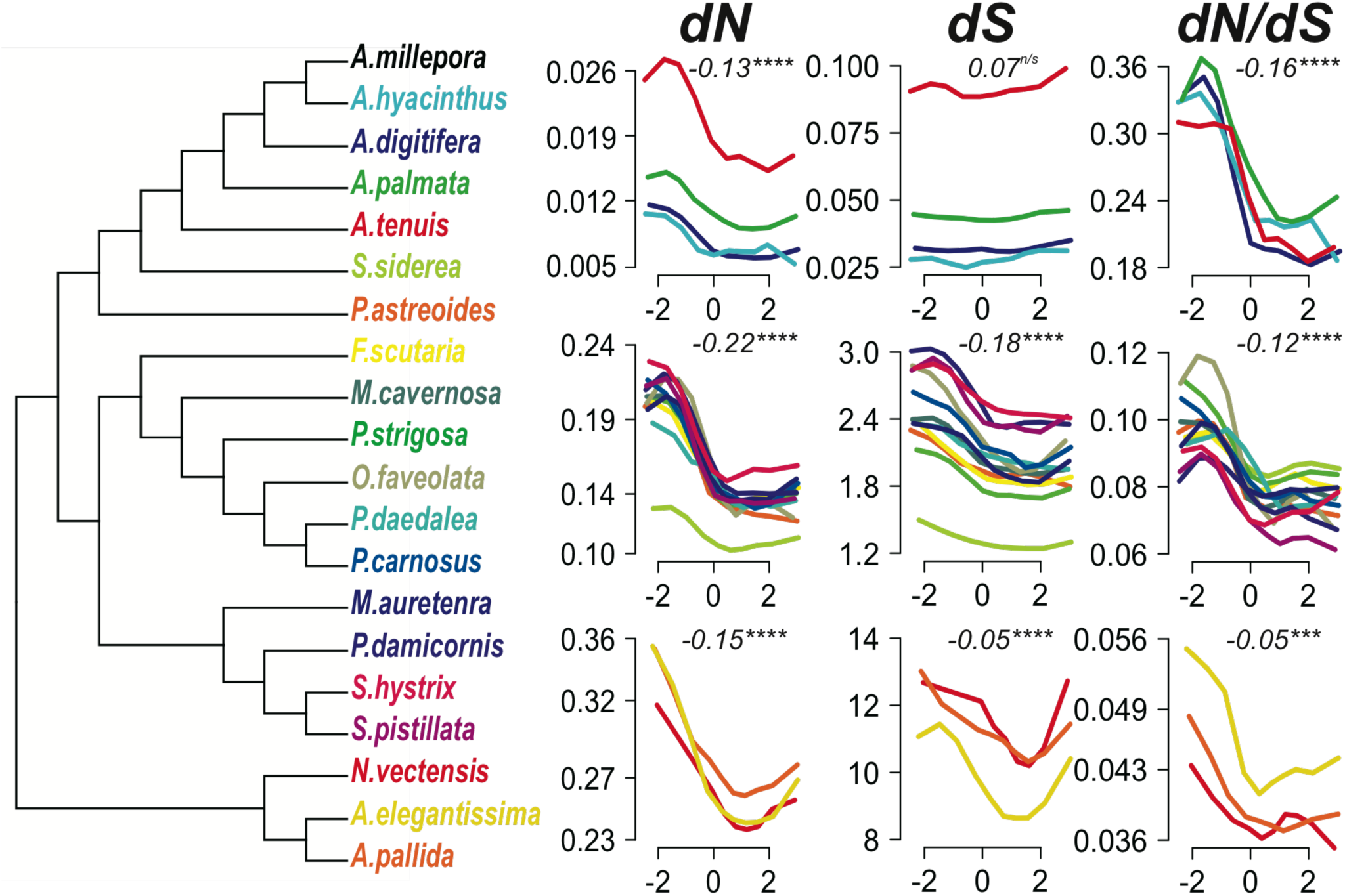
Relationship between MBD-score and substitution rates across the anthozoan phylogeny. All nodes in the phylogeny have 100% bootstrap support based on 1000 replicates. Line plots trace the mean substitution rates for all genes divided into ten quantiles ranked by MBD-score. Line color indicates which species *A. millepora* was compared with to estimate pair-wise substation rates. The top row of line plots show comparisons within *Acropora*. The middle row shows corals outside of *Acropora*. The third row shows comparisons with anemone species. Correlations were tested based on all datapoint using Spearman’s rho. Correlation and significance given for each panel indicate the median values for the included species. Significance notation: ns > 0.05; * < 0.05; ** < 0.01; ***< 0.001; **** < 0.0001.

The relationship between MBD-score and synonymous substitution rate (dS) was less pronounced than for dN, and varied with evolutionary proximity between species. Comparison of orthologs between *A. millepora* and other *Acropora* species showed no relationship between MBD-score and synonymous substitution rate (fig. 3). Comparisons with corals outside of *Acropora* however, showed a significant negative relationship, with an average of 17% higher mean dS for weakly methylated genes (fig. 3; supplementary fig. S6). The correlations with the three anemone species were weaker, although still significant. As most of these comparisons were saturated for synonymous substitutions they should be treated with caution. Analysis of dN/dS values gave similar results to dN for all groups of species (fig. 3).

### Strongly methylated genes show greater codon bias

Because DNA methylation alters mutation patterns, we hypothesized that gbM influences synonymous codon usage in stony corals. Specifically we predicted that strong gbM produces codon bias via mutational replacement of codons bearing CpG dinucleotides. To test this we correlated MBD-scores with three distinct indices of codon bias: frequency of optimal codons (Fop)(Ikemura 1981), codon adaptation index (CAI)(Sharp & Li 1987a), and effective number of codons (Nc)(Wright 1990). Fop and CAI each quantify preference for a set of optimal codons in the coding sequence. Higher values for these metrics indicate stronger codon bias. Nc quantifies nonrandom synonymous codon usage without assuming optimal codons. It is bounded between 1 (indicating complete bias, or use of only 20 codons for the 20 amino acids) and 64 (indicating completely neutral codon usage)(Wright 1990). All three indices correlated significantly with MBD-score (fig. 4). To assess the extent to which codon bias was driven by CpG hypermutability we recalculated CAI estimates using the same relative adaptiveness values ( *W* see methods) for each codon, but excluding the five amino acids coded for by CpG bearing codons (Serine, Proline, Threonine, Alanine and Arginine). This substantially weakened the correlation from 0.38 (Spearman’s rho; p << 0.0001) to 0.16 (Spearman’s rho), although it remained significant (p << 0.0001). In contrast, recalculation of CAI based solely on these five amino acids strengthened the correlation (rho = 0.42; supplementary fig. S7), indicating that hypermutability of CpGs due to gbM has a strong influence on codon usage.

**Figure 4:**
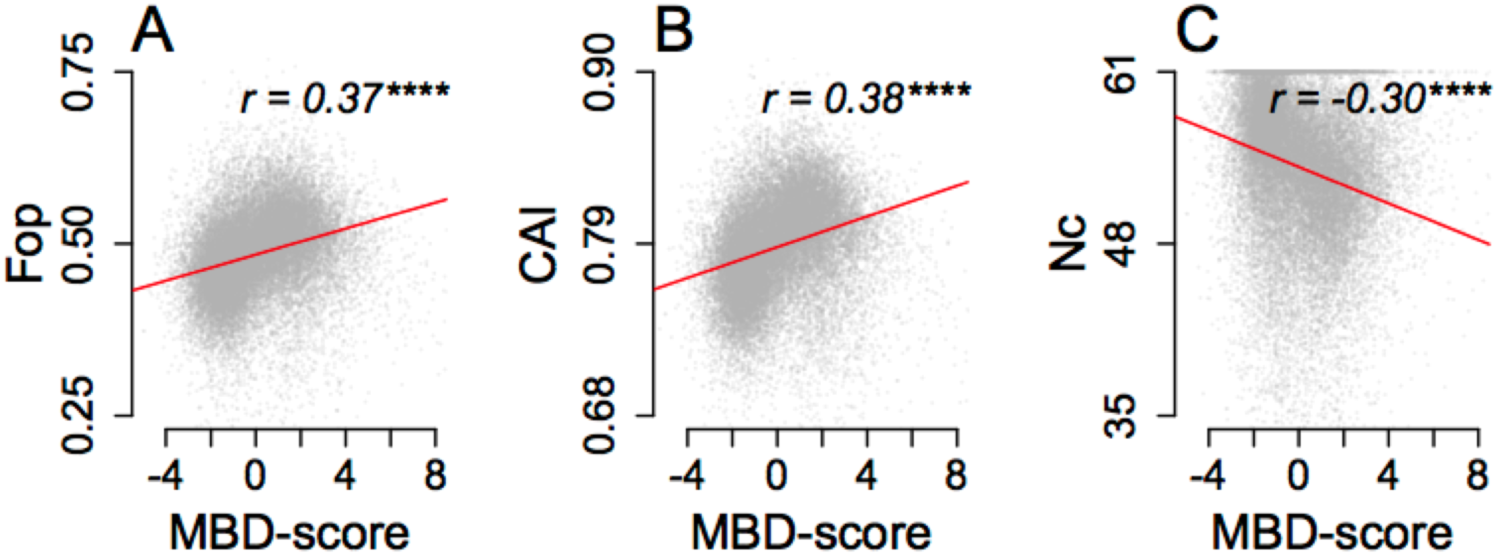
Correlation between MBD-score and indices of codon bias. (A) Frequency of optimal codons (Fop). (B) Codon adaptation index (CAI). (C) Effective number of codons (Nc). Correlation significance is based on Spearman’s rho (r). Red lines trace least squared linear regression.

### CpG codons are underrepresented in highly expressed genes

To further explore the influence of 5mC hypermutability on codon bias, we examined usage of CpG codons in highly expressed genes. As we did not have gene expression data for all species we first examined usage in annotated ribosomal protein genes with the assumption that these genes are highly expressed. For each species, relative synonymous codon usage (RSCU) of CpG codons was depressed in ribosomal protein genes (supplementary fig. S8A). To ensure that this did not result from variation in overall GC content we showed that mean RSCU of CpG codons was significantly lower than that of codons with GC, GG, or CC dinucleotides (t-tests; p for all species < 0.01).

For *A. millepora*, we assessed depression of CpG codons in highly expressed genes using three additional metrics: ΔRSCU (the difference in relative usage between the top 5% and bottom 5% expressed genes). rRSCU (the relative synonymous codon usage calculated for a concatenation of all ribosomal protein genes), and *W* (the relative adaptiveness of each codon; see methods). With one exception that had neutral usage, all CpG codons were underrepresented for all three metrics (supplementary fig. S8 B-D). Hence CpG bearing codons are depressed in highly expressed genes.

### Underrepresentation of CpG codons occurs through silent substitutions

To further illustrate that loss of CpG codons is due to 5mC hypermutability, we examined RSCU for the five amino acids coded for by CpG bearing codons. Four of these, (Threonine, Proline, Alanine and Serine), are coded for by NCG codons, in which the CpG occupies the second and third positions of the codon. For these codons, 5mC>T mutations on the sense strand necessarily produce amino acid changes, which are expected to be rare due to purifying selection. In contrast, 5mC>T substitutions of the corresponding CpG on the antisense strand produce silent substitutions (G>A within the codon)(supplemental fig.S9A). For this reason, we predicted that 5mC hypermutability would increase the usage of NCA codons at the expense of NCG codons. Plots of RSCU for synonymous codons against MBD-score illustrated positive relationships for NCA codons (Spearman’s rho between 0.156 and 0.196; p << 0.001) and opposing negative relationships for NCG codons (fig. 5). Correlations of NCA codon usage with MBD-score were significantly stronger than for other synonymous codons (t-test; p < 0.01). Moreover, all NCA codons were identified as optimal codons (supplemental table S1). and their mean relative adaptiveness (for which the maximum is 1) was 0.99 (supplemental table S2). Together these data indicate that that NCA codons replace NCG codons in strongly methylated genes.

**Figure 5:**
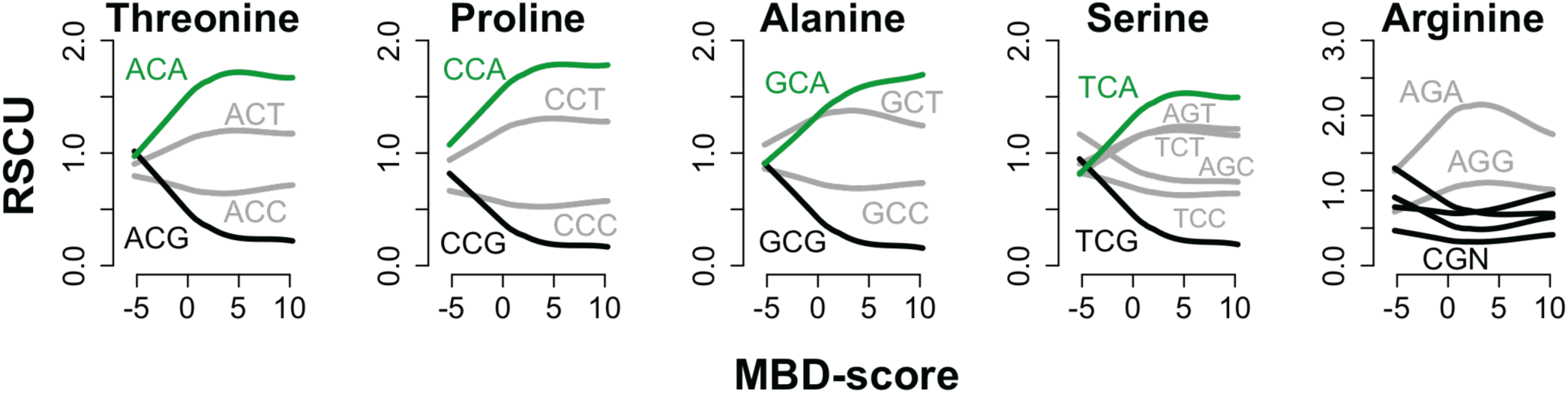
Depression of CpG bearing codons occurs via replacement with synonymous NCA codons. Lines show smoothed traces of the relationship between Relative Synonymous Codon Usage (RSCU) and MBD-score for the indicated codon. Black lines indicate CpG bearing codons. Green lines indicate NCA codons. Grey lines indicate all other codons. For NCA codons, correlations were between RSCU and MBD-score were between 0.156 and 0.196 (Spearman’s rho; p << 0.001). Opposing trends for NCA and NCG codons support the inference that NCA codons replace NCG codons in strongly methylated genes.

The second group of CpG bearing codons is the CGN codons, which code for arginine. These are expected to evolve differently because 5mC>T substitutions on both the sense and antisense stands produce amino acid changes (supplemental fig.S9A). Although the trend is weak (r = −0.06; p < 0.0001), arginine content is negatively correlated with MBD-score (supplemental fig.S9B), suggesting a slight shift in mutation-selection balance for arginine in strongly methylated genes.

### Summarizing interrelationships between gene characteristics

To summarize the relationships between gbM and other gene characteristics we performed principal component analysis (PCA) on all coding regions for which we had MBD-scores and substitution rate estimates. Pair-wise estimates of dN and dS between *A. millepora* and *Siderastrea siderea* were used because it was the species outside of the genus *Acropora* with the greatest number of orthologs. Substitution rates based on other species produced qualitatively similar results. Variation in measures of gbM and codon bias was captured largely by the first principal component (34.0% variance explained) (fig. 6). While the indices of codon bias often correlated most strongly with one another, the strongest alternative predictor for all three was CpGo/e (supplemental table S3). Variation in transcript abundance, gene length, and substitution rates was captured largely by the second principal component (14.2% variance explained)(fig. 6).

**Figure 6:**
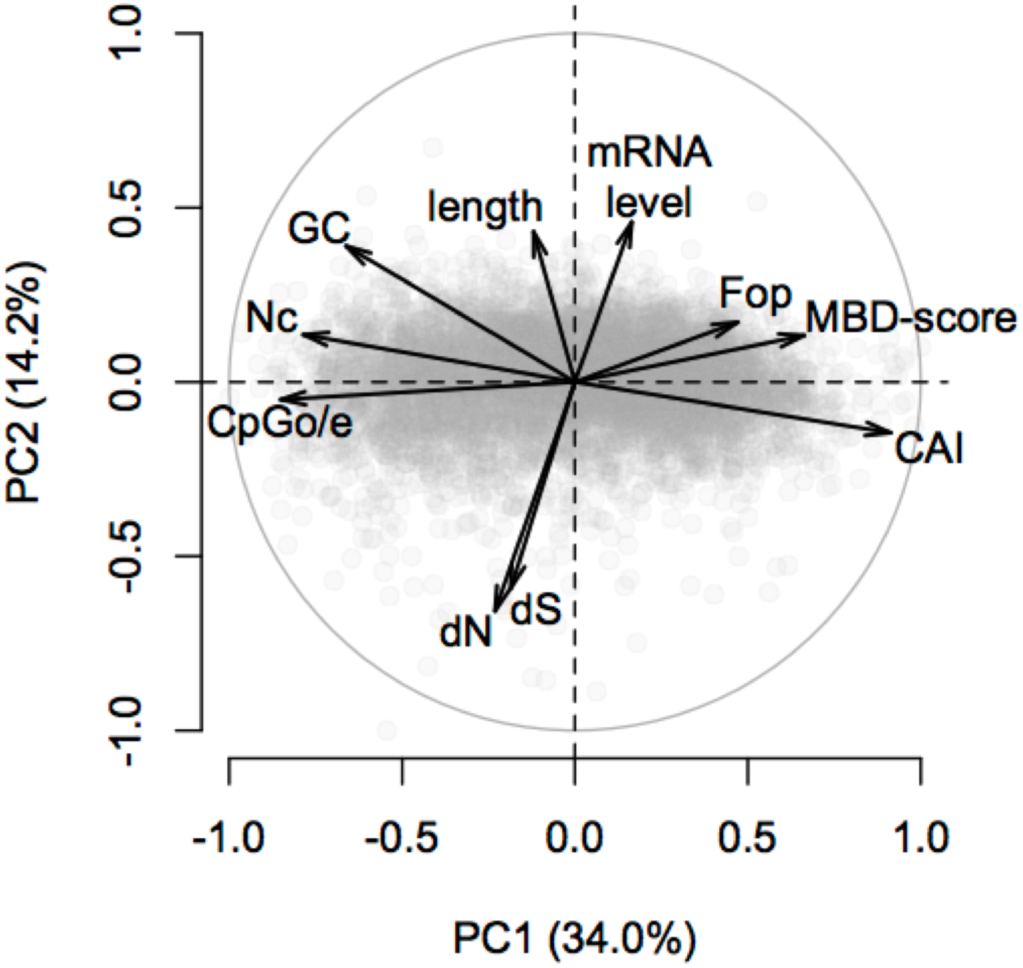
Principal component analysis of gene features in *A. millepora*. The first principal component explained 34.0% of variation and correlated primarily with measures of gbM and codon bias. The second principal component explained 14.2% of variation and correlated primarily with gene length, transcript abundance, and substitution rates. Variables included in the analyses are: normalized CpG content (CpGo/e), effective number of codons (Nc), GC content of coding regions (GC content), nonsynonymous substitution rate (dN), synonymous substitution rate (dS), length of coding region (length), transcript abundance (mRNA level), Frequency of optimal codons (Fop), log2 fold difference between captured and flowthrough fractions of methylation binding domain enrichment libraries (MBD-score), and codon adaptation index (CAI), Substitution rates are pairwise estimates between *A. millepora* and *S. siderea*.

## Discussion

### Gene body methylation is a signature of broad and stable expression

We showed that strongly methylated genes tend to have constitutive and ubiquitous functions and are less likely to be differentially expressed across developmental stages and environmental regimes. These results confirm earlier findings from diverse metazoans including a mollusk (Gavery & Roberts 2010; Gavery and Roberts 2014), arthropods (Elango *et al*. 2009; Wang *et al*. 2013), and cnidarians (Sarda *et al*. 2012; Dimond & Roberts 2015) as well as *Arabidopsis* (Aceituno *et al*. 2008; Takuno & Gaut 2012), The relationship with differential expression in response to environmental regimes suggests the intriguing possibility that gbM could modulate gene expression plasticity.

### Gene body methylation and evolutionary rates

Using pair-wise comparisons between *A. millepora* and 19 other anthozoan species we show that gbM negatively correlates with substitution rates. This finding is consistent with previous results from invertebrates (Park *et al*. 2011; Sarda *et al*. 2012), mammals (Chuang and Chiang 2014) and plants (Takuno & Gaut 2012). Still, principal component analysis revealed that while substitution rates are negatively correlated with gbM, they correlate more strongly with transcript abundance, as has been previously shown in bacteria, plants, fungi, and animals (Drummond & Wilke 2008; Yang & Gaut 2011). The ubiquitous negative correlation between substitution rate and transcript abundance is explained by stronger purifying selection against protein misfolding in highly expressed genes. Because highly expressed proteins have a greater cumulative opportunity for misfolding, their mutations pose greater fitness costs than those in lowly expressed genes (Drummond *et al*. 2005). Similar logic can be applied to broadly expressed genes, as they are active in a greater number of cells and tissues (Duret & Mouchiroud 2000) and undergo more translational events at the scale of the entire organism. Whereas non-synonymous substitutions affect the probability of protein misfolding through direct destabilization, synonymous substitutions most likely exert a similar but weaker effect by lowering translational accuracy (Akashi 1994; Drummond and Wilke 2008). Hence both dN and dS are expected to be lower in highly and broadly expressed genes. We have shown that in our system, highly expressed genes tend to be strongly methylated (supplemental fig. S3), and that strongly methylated genes tend toward broad, constitutive transcription (fig 2; supplemental fig. S2). We conclude that the observed correlation between gbM and substitution rates is most parsimoniously explained by the occurrence of gbM on genes that are under stronger purifying selection because of their expression patterns.

### Gene body methylation shapes codon usage

Codon bias occurs for two reasons. One mechanism is mutational bias, where differences in mutation rates across species and genomic contexts produce non-random variation in synonymous codon usage (Plotkin & Kudla 2011). The second mechanism is natural selection, which requires that synonymous mutations affect organismal fitness (Behura & Severson 2013). We found that gbM correlates strongly with three separate indices of codon bias (fig. 4). This appears to be due largely to mutational patterns. CpG codons were depressed in highly expressed genes (supplemental fig.S8), and the strongest predictor of codon bias was historical germline methylation as measured by CpGo/e (fig. 6; supplementary table S1). Analysis of RSCU values for NCG codons was consistent with codon bias arising largely through silent 5mC>T substitutions on the antisense stand (fig. 5; supplemental fig.S9A). In other words, gbM causes a shift in usage of NCG codons to NCA codons.

It should be noted that CAI correlated significantly with gbM even when amino acids with CpG codons were excluded from the analysis, (supplemental fig.S7). This suggests that an additional mechanism to CpG mutation, potentially natural selection, increases codon bias within strongly methylated genes. When assessing whether codon bias is due to selection, researchers examine whether it occurs in genes for which translation accuracy and efficiency are most important. Evidence that codon bias is due to selection includes: 1) positive correlation with expression level, 2) positive correlation with breadth of expression, and 3) negative correlation with synonymous substitution rate (Sharp & Li 1987b; Duret 2002; Plotkin & Kudla 2011). As we have shown, gbM in *A. millepora* covaries with each of these factors.

It is interesting that gbM is not only a potent source of codon bias, but also occurs preferentially on genes for which adaptive codon usage is most important. We propose that the systematic codon bias produced by gbM could itself be adaptive, establishing an equilibrium in which particular sets of preferred and unpreferred codons are maintained in constitutively active genes. Optimal translation dynamics could then be achieved through evolution of tRNA abundances to match these preferred codons, obviating the need for selection of individual codons on a site-by-site basis. To put it another way, selection coefficients for individual synonymous codons will be exceptionally small (Bulmer 1987). In contrast, if a set of preferred codons is mutationally established in constitutively expressed genes, alleles that control the abundance of appropriate tRNAs would have stronger fitness effects more amenable to natural selection. To be clear, we are not proposing that gbM originally evolved for this purpose. However, if its original function was linked with constitutively active genes, as appears to be the case from studies of plants (Coleman-Derr and Zilberman 2012; Takuno & Gaut 2012), invertebrates (This study) and mammals (Baubec *et al*. 2015), then CpG depression coupled with coevolution of tRNAs provides an efficient means of maintaining optimal codon usage in the genes where it is most beneficial. This hypothesis could be tested through phylogenetic comparison of tRNA abundances between clades that have independently lost or retained gbM.

An advantage of mutation-driven codon bias is that it could be maintained even in the absence of efficient selection, so that it would be most beneficial for organisms with relatively small population sizes. If the adaptive value of gbM is indeed related to maintenance of codon usage, it is not surprising that organisms such as yeast, fly and worm are able to exist without it: due to their large population sizes their optimal codon usage can be maintained by selection alone.

### Conclusions and outlook

Here we present three primary findings on gene body methylation in stony corals: 1) gbM is most pronounced in genes with broad and stable expression across time and space; 2) gbM predicts slow sequence evolution 3) hypermutability due to gbM is a principal driver of codon bias. What does this tell us of the ancestral function of gbM? Preferential occurrence on broadly and actively expressed genes in plants (Coleman-Derr and Zilberman 2012; Takuno & Gaut 2012) and the basal metazoan examined here indicates an evolutionarily ancient function involving selective pressure for accurate and stable gene expression. One means of improving translation fidelity is the use of optimal codons. If codon bias introduced by gbM corresponds to tRNA abundance, it could be a previously unrecognized driver of adaptive codon evolution.

## Materials and methods

### Sequence Data and Computational Tools

Transcriptomic data from 17 species of Scleractinia (stony corals) and 3 species of Actiniaria (anemones) were downloaded from the web (Supplementary table S4; Schwarz *et al*. 2008; Sunagawa *et al*. 2009; Polato *et al*. 2011; Shinzato *et al*. 2011; Moya *et al*. 2012; Kenkel *et al*. 2013; Lubinski & Granger 2013; Sun *et al*. 2013; Maor-Landaw *et al*. 2014; Nordberg *et al*. 2014; Willette *et al*. 2014; Kitchen *et al*. 2015; Davies *et al*. forthcoming). Instructions, scripts, and example output files for computational methods used in this study are available on GitHub (https://github.com/grovesdixon/metaTranscriptomes). Gene Ontology and KOG annotations were applied as described in (Dixon *et al*. 2015). Instructions and scripts for the gene annotation pipeline are available on GitHub (https://github.com/z0on/annotatingTranscriptomes). Significance for enrichment of KOG terms across MBD-scores was tested using Mann-Whitney U tests implemented in the R package KOGMWU as in Dixon et al. (2015).

### Ortholog Identification and alignment

Orthologs were identified based on reciprocal best Blast hits between extracted protein sequences. First, protein coding sequences were extracted for each transcriptome based on alignments (e-value cutoff = 1e-5) to a reference proteome using BlastX (Altschul *et al*. 1997) and a custom Perl script CDS_extractor_v2.pl (https://github.com/z0on/annotatingTranscriptomes) that identifies and corrects frame shift mutations within the BlastX-aligned sequences. The reference proteome was a concatenation of the *Nematostella vectensis* (Nordberg *et al*. 2014) and *Acropora digitifera* (Shinzato *et al*. 2011) reference proteomes. The extracted coding sequences were then translated to produce a protein sequences. The protein sequences for all pairs of species were reciprocally blasted using BlastP (Altschul *et al*. 1990). Because our MBD-seq dataset was generated from *A. millepora*, we used its sequences as anchors for orthologous groups. First an initial set of candidate orthologs was compiled based on reciprocal best hits with *A. millepora*. Only hits with alignment lengths >75% of the subject sequence and an e-value < 1e-5 were retained. This initial set was then refined to include only sequences that were reciprocal best hits with >= 50% of other candidate orthologs within the group (supplementary fig. S10). Orthologous groups with fewer than three (15%) representative species were excluded. For building the species tree, a separate, highly conserved set of orthologs was assembled with amino acid identity > 75%. These were further filtered by retaining only orthologs with representative sequences from > 80% of species. As a final filter, we used cluster analysis of dS values to identify likely paralogs and spurious orthologs. For each species a three component Gaussian mixture model was fit to the pairwise dS estimates with *A. millepora*. The first two components were assumed to capture the true orthologs, the third component was assumed to have captured paralogs and false positive orthologs (supplemental fig. S11). Mean dS for the third component was on average 60 times higher than the second highest component. On average 10% of ortholog calls were flagged as false positives and removed. Amino acid sequences for each ortholog were aligned with MAFFT (Katoh & Standley 2013) using the ‘localpair’ algorithm. The protein alignments were then reverse translated into codon sequences using Pal2Nal (Suyama *et al*. 2006).

### Substitution rate analyses

To estimate substitution rates (dS and dN) we used codeml in the PAML software package (Yang 2007). Substitution rates were estimated using pair-wise comparisons between all pairs of taxa that had representative sequences for each ortholog. Example codeml control files for the pair-wise comparisons are available on GitHub (https://github.com/grovesdixon/metaTranscriptomes).

### Building species tree

Based on a set of highly conserved ortholog sequences we constructed a species tree using RAxML (Stamatakis 2014). The rapid bootstrapping algorithm was run using the GTRGAMMA model and 1000 iterations. We decided to use putative orthologs with representative sequences in > 80% of taxa through iterations of tree building. The best trees from ortholog sets using 40%, 50% and 60% cutoffs all gave the same topology. The best tree using the 80% cutoff was chosen because it had highest bipartition bootstrap values.

### Library preparation for MBD-seq

To quantify gbM in *Acropora millepora* we used methyl-CpG binding domain protein-enriched sequencing (MBD-seq). Enrichment reactions were performed using the MethylCap kit (Diagenode Cat. No. C02020010). Seven enrichment reactions were performed. From each reaction both the captured fraction (strongly methylated) and flow-through (weakly methylated) was retained for sequencing. Adapter ligation using a NEBnext kit (New England Biolabs®), library quality assessment using a Bioanalyzer (Agilent Technologies), and sequencing on a HiSeq 2500 platform (Illumina®) were performed by the University of Texas Genome Sequencing and Analysis Facility. Further details on library preparation are given in the supplemental materials (supplemental methods).

### Analysis of gene body methylation

Gene body methylation was quantified as the fold difference in coverage between the MBD-enriched and flow-through libraries. Raw reads from the MBD-sequencing libraries were trimmed using cutadapt (Martin 2011) and quality filtered using Fastx toolkit (http://cancan.cshl.edu/labmembers/gordon/fastx_toolkit/). Reads were then aligned to coding sequences extracted from the *A. millepora* reference transcriptome (Moya *et al*. 2012) as described above. DESeq2 (Love *et al*. 2014) was used to calculate the log2 fold difference between the MBD-enriched and flow-through libraries. We used this log2 fold difference, which we refer to as MBD-score, as our quantification of the strength of gbM for each gene. Negative values indicate weak methylation and positive values indicate strong methylation. To examine the distribution of MBD-scores we used the R package Mclust (Fraley & Raftery 2007). We first assessed the optimal mixture model and number of components based on Bayesian Information Criterion (BIC). The optimal number of components was greater than one with little change in BIC beyond two components (supplemental fig. S1). Based on this result we fitted a two-component mixture model to the MBD-scores (supplemental fig. S1).

Because of the hypermutability of 5mC, genes that are strongly methylated in the germline become deficient in CpG dinucleotides over evolutionary time (Sved & Bird 1990). As a result, normalized CpG content (CpGo/e) can be used to estimate historical germline methylation. This metric has been shown to correlate closely with direct measures of gbM (Zemach *et al*. 2010; Sarda *et al*. 2012). To corroborate that our measure of gbM also correlated with CpGo/e we calculated it for the *A. millepora* coding regions as described in Dixon et al. (2014). To control for effects on gene length, CpGo/e was calculated based on the first 1000 bases of each sequence.

### Gene expression Datasets

To test for correlations between MBD-score and transcriptional variation we used gene expression data from two previous experiments. Both datasets were generated using Tag-based RNA-seq (Meyer *et al*. 2011) from samples of *A. millepora* taken from the central Great Barrier Reef, Australia. The current laboratory and bioinformatics protocols for analysis of Tag-based RNA-seq are available on GitHub (https://github.com/z0on/tag-based_RNAseq). The first dataset, described in (Dixon *et al*. 2015), included 12 adult samples (six genotypes from Princess Charlotte Bay and six from Orpheus Island: Great Barrier Reef Marine Park Authority permit G38062.1) and 30 offspring larval samples. Adults were sampled following acclimatization period of 5 days at 28°C in a common shaded raceway. Larvae were sampled five days postfertilization (advanced planula stage), reared at 28°C. Variation in gene expression between adults and larvae was analyzed using DESeq2 (Love *et al*. 2014). Comparisons between MBD-score and transcript abundance were based on counts from adult samples transformed to a log2 scale using the rlogTransformation function. Mean expression levels from this dataset were also used to calculate indices of codon bias described below. The second dataset described in (Dixon *et al*. 2014) included 56 colony fragments reciprocally transplanted between two environmentally distinct reefs: Keppel and Orpheus Island (Keppel: 23°09S 150°54E and Orpheus 18°37S 146°29E: Great Barrier Reef Marine Park Authority permit G09/29894.1). Expression profiles from these samples were analyzed with respect to the transplantation site to examine variation in gene expression due to environmental conditions.

### Codon Bias

We tested for relationships between MBD-score and synonymous codon usage using four metrics: relative synonymous codon usage (RSCU)(Sharp *et al*. 1986), frequency of optimal codons (Fop)(Ikemura 1981; Behura & Severson 2013), codon adaptation index (CAI)(Sharp & Li 1987a), and the effective number of codons (Nc)(Wright 1990). Fop and Nc were estimated using the CodonW (Peden 1999; http://codonw.sourceforge.net//culong.html). CAI was estimated using custom python scripts. Further details on estimation of codon bias are given in the supplemental material (supplementary methods).

### Statistical Analyses

Statistical analyses of the relationship between MBD-score and other gene characteristics were performed using R (R Core Team 2015). Significance for correlations was tested using Spearman’s *rho*. Significance tests for differences in counts between the strongly methylated and weakly methylated classes were performed using Fisher’s exact tests (Fisher 1922). Principal component analysis was performed using prcomp function in R.

## Acknowledgements

This work was supported by the National Science Foundation (grant DEB-1054766 to M.V.M).

